# A Human Genetics Framework for De-risking Gene Editing Targets for Hematopoietic Cell and Gene Therapy

**DOI:** 10.64898/2025.12.15.694499

**Authors:** Abhinav Dhall, Alyssa Pyclik, Matt Ung, Julia Etchin, Yonina Keschner, Huanying Gary Ge, Tirtha Chakraborty, John R. Lydeard

## Abstract

Developing novel therapeutics requires robust early-stage target de-risking to ensure safety and efficacy. We developed a scalable proteogenomic framework integrating population-scale human genetics and plasma proteomics to identify genes tolerant of inactivation (i.e., dispensable) within hematopoietic compartments, thereby enabling safer targeted immunotherapies. Using *CD33* as a validated benchmark, we observed that naturally occurring loss-of-function (LoF) variants lead to concordant RNA and protein depletion, supporting functional gene inactivation. Early clinical results from the Trem-Cel trial (NCT05945849) further provide proof of concept that deletion of dispensable antigens can enable safe and effective immunotherapy in humans. We extended this approach genome-wide in the UK Biobank and identified 237 candidate dispensable genes, filtered by plasma proteomic data and hematopoietic expression, highlighting *LY75* (CD205) as a novel candidate with strong proteogenomic evidence of LoF tolerance. This work establishes a generalizable, quantitative proteogenomic framework for systematic prioritization of dispensable gene targets for editing, providing a foundation for next-generation cell and gene therapies that minimize on-target, off-tumor toxicities.

## Introduction

Fewer than 10% of candidates that enter Phase 1 clinical trials ultimately receive FDA approval, underscoring the critical need for early-stage approaches to assess safety and efficacy.^1^ Human genetic evidence has emerged as a powerful tool for de-risking drug targets. In a large analysis encompassing approximately 23,000 drugs, those supported by human genetic evidence were approximately twice as likely to obtain regulatory approval.^2,3^ Naturally occurring loss-of-function (LoF) variants represent “experiments of nature” that reveal which genes can be disrupted without major adverse consequences. One such precedent is *PCSK9*, where naturally occurring LoF variants guided the development of safe and effective inhibitors.^4^

This concept is especially relevant for gene-edited cell-based applications, where therapeutic gene ablation introduces unique safety considerations. Identifying genes that are genetically dispensable in humans offers a path to safer targets for therapeutic editing, particularly within hematopoietic compartments, where cell and gene-based therapies are most advanced. In acute myeloid leukemia (AML), current immunotherapies including antibody-drug conjugates, bispecific T-cell engagers, and chimeric antigen receptor (CAR) T-cell therapies target antigens like CD33 and CD123. While these antigens are highly expressed on leukemic blasts, they are also present on normal hematopoietic cells, leading to on-target, off-tumor toxicities including prolonged cytopenias and other clinical complications.^5^ Gene editing offers a potential solution by selectively deleting dispensable target antigens in healthy donor-derived hematopoietic stem and progenitor cells (HSPCs) prior to transplantation.^6,7^ After transplantation, immunotherapy directed against the deleted antigen, still present on unedited AML blasts, can selectively eliminate residual leukemic cells while sparing the gene-edited donor cells. This strategy entered clinical investigation with *CD33*-null HSPCs (NCT04849910, NCT05945849, NCT05662904). Notably, early clinical results from the Trem-Cel trial (NCT05945849) demonstrated rapid primary hematopoietic engraftment with durable CD33-negative hematopoiesis. Importantly, we also observed protection from CD33-directed therapy, providing proof of concept for targeting genetically dispensable antigens in the clinic.^8,9^

Gene knockout approaches are also being explored to enhance T cell-based therapies. For example, deletion of *CD5* or *CD7* reduces CAR T cell fratricide, tonic signaling, and early exhaustion, thereby enhancing anti-tumor activity.^10,11,12,13^ Although these strategies are advancing to clinical trials (NCT06534060, NCT06420089), most lack validation from population-scale human genetic data, leaving uncertainty around target dispensability.

To address this gap, we developed a framework that integrates human genetic variation with plasma proteomics to systematically evaluate gene dispensability. Using well-curated resources such as the Genome Aggregation Database (gnomAD) and the UK Biobank (UKB), we combined gene constraint metrics, LoF variant data, and protein abundance measurements to quantitatively assess tolerance to gene disruption. Gene constraint was evaluated using probability of loss-of-function intolerance (pLI) and loss-of-function observed/expected upper bound fraction (LOEUF).^14^ Predicted LoF variant data included frameshift, essential-splice, start-loss and stop-gain alleles, referred to as protein-truncating variants (PTVs). We evaluated the former as well as all missense variants using protein abundance from plasma proteomic measurements available in the UKB.^15^

For this study, we operationally define dispensability as the presence of high-confidence homozygous LoF variants observed in population cohorts, together with evidence of transcript and/or protein depletion in carriers and the absence of overt developmental or clinical consequences in UKB participants. This definition provides a consistent standard for evaluating whether a gene can be targeted for therapeutic knockout. We first illustrate this framework using *CD33* as a benchmark and contrast these findings with genes that, while not strictly essential for cell survival, exhibit strong evolutionary constraint and clinically meaningful loss-of-function phenotypes, such as CD45 (*PTPRC*). We then extended this analysis genome-wide within the UKB to identify additional candidate genes.

To ensure translational relevance, we focused on genes expressed within hematopoietic compartments where therapeutic gene-edited strategies are most directly applicable. This analysis encompassed all protein-coding genes in the 470K UK Biobank exome dataset, which were subsequently filtered using plasma proteomic data and hematopoietic expression profiles to prioritize genes most relevant for therapeutic gene editing. Among these, *LY75* emerged as the top-ranked candidate, with strong proteogenomic evidence of tolerance to LoF and potential utility as a therapeutic knockout target.

## RESULTS

### CD33 Defines Quantitative Parameters of Gene Dispensability

*CD33* is a key example of a dispensable gene, supported by genetic, experimental, and clinical evidence.^6,7,16^ *CD33* shows minimal evolutionary constraint (LOEUF ≈ 1.07, pLI = 0 in gnomAD v4.1.0), consistent with tolerance to LoF. In gnomAD v4.1.0, 612 individuals out of 730,947 exomes and 76,215 genomes sequenced, harbor homozygous LoF variants in *CD33*, most commonly (94%) a four–base pair deletion in exon 3 (rs201074739; 19:51729103:CCCGG:C) that introduces a premature stop codon resulting in loss of *CD33* expression. The age distribution of homozygous carriers was similar to that of heterozygotes and noncarriers, with several individuals reaching advanced ages (64–93 years), further supporting the absence of detrimental effects. Together, these data exemplify the parameters by which we define dispensability: the presence of homozygous LoF alleles, molecular evidence of gene inactivation, and evidence that gene disruption does not impair development or longevity.

Experimental studies demonstrate that *CD33* deletion does not compromise human HSPC function.^6,7,8^ CD33-deficient human HSPCs demonstrated normal engraftment and differentiation in immunocompromised mice, while providing robust protection from the cytotoxic effects of CD33-directed companion therapeutics. Additionally, autologous *CD33* knockout HSPC transplantation in non-human primates supported long-term multilineage engraftment of gene-edited cells that maintained normal hematopoietic function.^6,7^ In humans, *CD33*-null HSPCs are expected to reconstitute a hematopoietic system resistant to anti-CD33 therapies such as Gemtuzumab ozogamicin (GO).^9^ Notably, early clinical results from the Trem-Cel trial (NCT05945849) demonstrated rapid primary hematopoietic engraftment with durable CD33-negative hematopoiesis and protection from GO-associated hematopoietic toxicity, providing proof of concept for targeting genetically dispensable antigens to enhance safety in the clinic.^8^ Together, these data establish *CD33* as a proof-of-concept benchmark for identifying genetic parameters associated with dispensable genes.

### CD33 LoF Variants Reduce mRNA Expression and Protein Abundance

Having established that genetic *CD33* deletion is well tolerated in humans and preclinical models, we next examined the molecular consequences of various naturally occurring LoF variants of *CD33*. Because annotated LoF variants do not always yield complete gene inactivation, we sought to define the molecular signatures of bona fide LoF by quantifying the effects of *CD33* LoF variants on both transcript and protein abundance. To do so, we analyzed allele-specific expression (ASE) in GTEx and plasma proteomics in the UK Biobank. Heterozygous carriers of the common *CD33* LoF variant (rs201074739; 19:51729103:CCCGG:C) showed reduced expression of the LoF variant to the reference allele, consistent with nonsense-mediated decay (Figure 1A).^17^

**Figure 1:**
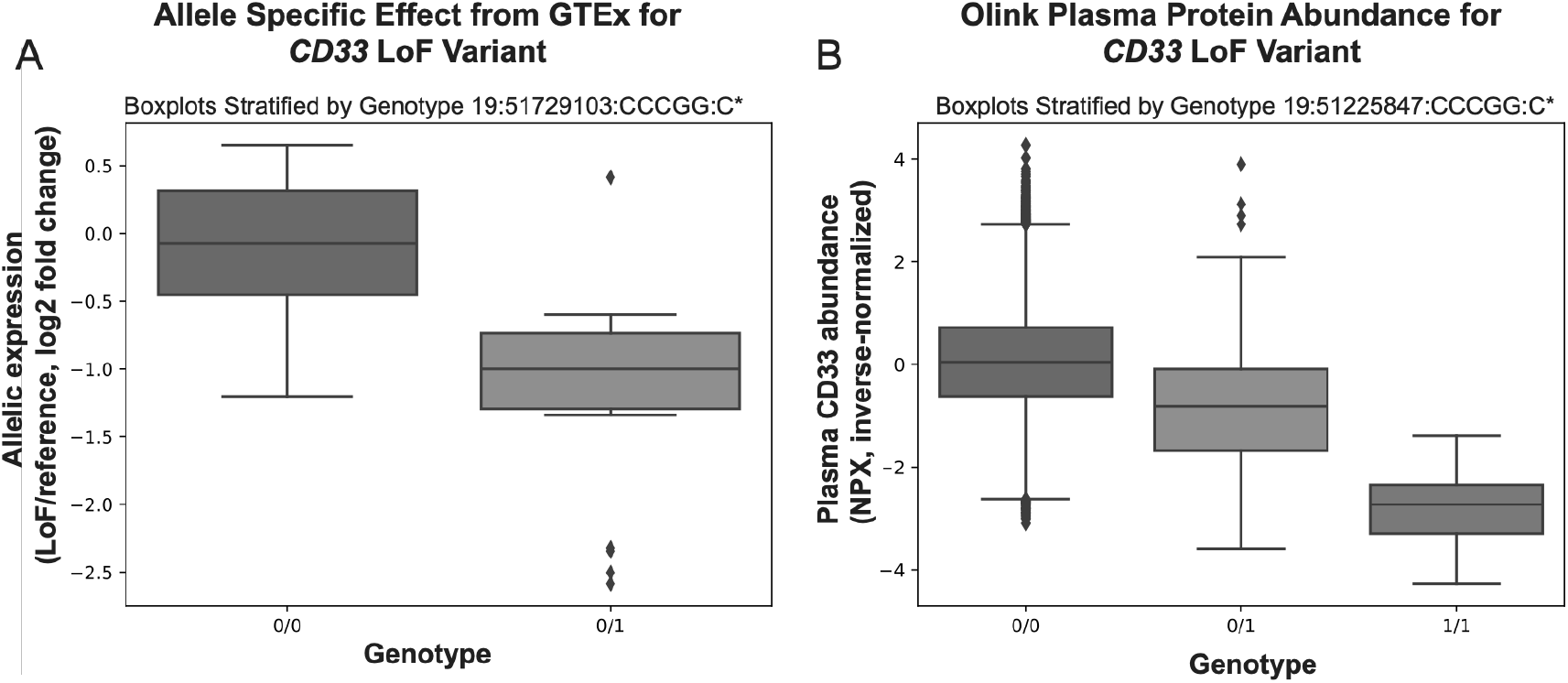
CD33 LoF variant reduces mRNA and plasma protein abundance. (A) Allele-specific expression of the common CD33 LoF variant in GTEx. Heterozygous carriers (0/1, n=524) exhibit reduced expression of the LoF allele relative to the reference (0/0). p < 1×10^−6^. (B) Plasma CD33 protein abundance (Olink NPX, inverse-normal transformed) in UKB participants stratified by genotype. A stepwise, genotype-dose reduction (0/0 > 0/1 > 1/1) in CD33 levels is observed (p < 1×10^−8^). Boxplots show median and interquartile range; points = individuals; line = mean ± 95% CI. 0= reference, 1-LoF allele. In UKB n=46,931 for 0/0, n=2452 for 0/1 and n=36 for 1/1. Note: Genome coordinates differ between GTEx and UKB due to different reference builds, but both refer to the CD33 rs201074739; the CCCGG:C LoF variant.

Protein measurements revealed a clear dose-dependent effect of genotype on CD33 abundance in plasma. Homozygous carriers (1/1) had the lowest CD33 abundance, heterozygotes (0/1) intermediate, and wild-type individuals (0/0) showed the highest (Figure 1B). These associations were highly significant (p < 1e-8). Median normalized protein expression (NPX, inverse-normal transformed) values were −2.1 Standard Deviations (SD) for homozygous carriers, −0.8 SD for heterozygotes, and 0.1 SD for wild-type individuals, illustrating the stepwise, genotype-dose relationship. These concordant reductions at both transcript and protein levels support functional inactivation of *CD33 in vivo* and establish it as a quantitative benchmark for gene dispensability. With this benchmark defined, we applied the same framework to systematically identify additional dispensable therapeutic targets. We next evaluated candidate antigens prioritized from prior studies and, using the genetic parameters of “dispensability” outlined in Sup Table 1, we performed a genome-wide variant assessment of LoF variants. To ensure translational relevance, we integrated UKB plasma proteomics data and restricted analysis to genes expressed within hematopoietic compartments (Figure 3A; Methods).

### Assessment of Candidate Hematopoietic Target Genes

To test whether our framework distinguishes functionally important from dispensable antigens, we next applied it to well-studied immunotherapy targets (Table 1). We expected that genes with strong evolutionary constraint-characterized by low LOEUF, high pLI, and known clinical null phenotypes would not meet our dispensability criteria (Sup Table 1), whereas genes more tolerant of inactivation would. *PTPRC* (CD45), a functionally important gene, served as a counterpoint to *CD33*, while *IL3RA* and *KIT* provided intermediate examples of tolerability (Table 1). These comparisons corroborate that our framework separates validated dispensable targets from functionally important genes whose loss produces clinically significant phenotypes. *PTPRC (CD45)* illustrates a gene under strong evolutionary constraint (LOEUF= 0.34, pLI=1). Clinically, biallelic CD45 LoF leads to misregulation of B and T cells resulting in severe combined immunodeficiency (SCID), which is fatal in childhood.^18,19^ Unlike CD33, CD45 has no confirmed homozygous null individuals in healthy populations, underscoring its functionally important role. Consistent with this, we found no replicated, homozygous nulls validated at the protein level in population cohorts. There is one homozygous LoF carrier annotated in gnomAD v4.1.0 with the variant 1:198729171:GGTAA:G predicted to result in the deletion of four nucleotides in the splice donor site of exon 16 of the canonical *PTPRC* mRNA transcript (ENST00000442510.8). However, this variant did not result in transcript or plasma protein depletion in the UKB genetics and Olink plasma protein datasets, respectively, suggesting it is not a true LoF. Further work will be needed to fully assess the molecular impact of the variant to determine if it is a true LoF variant.

**Table 1:**
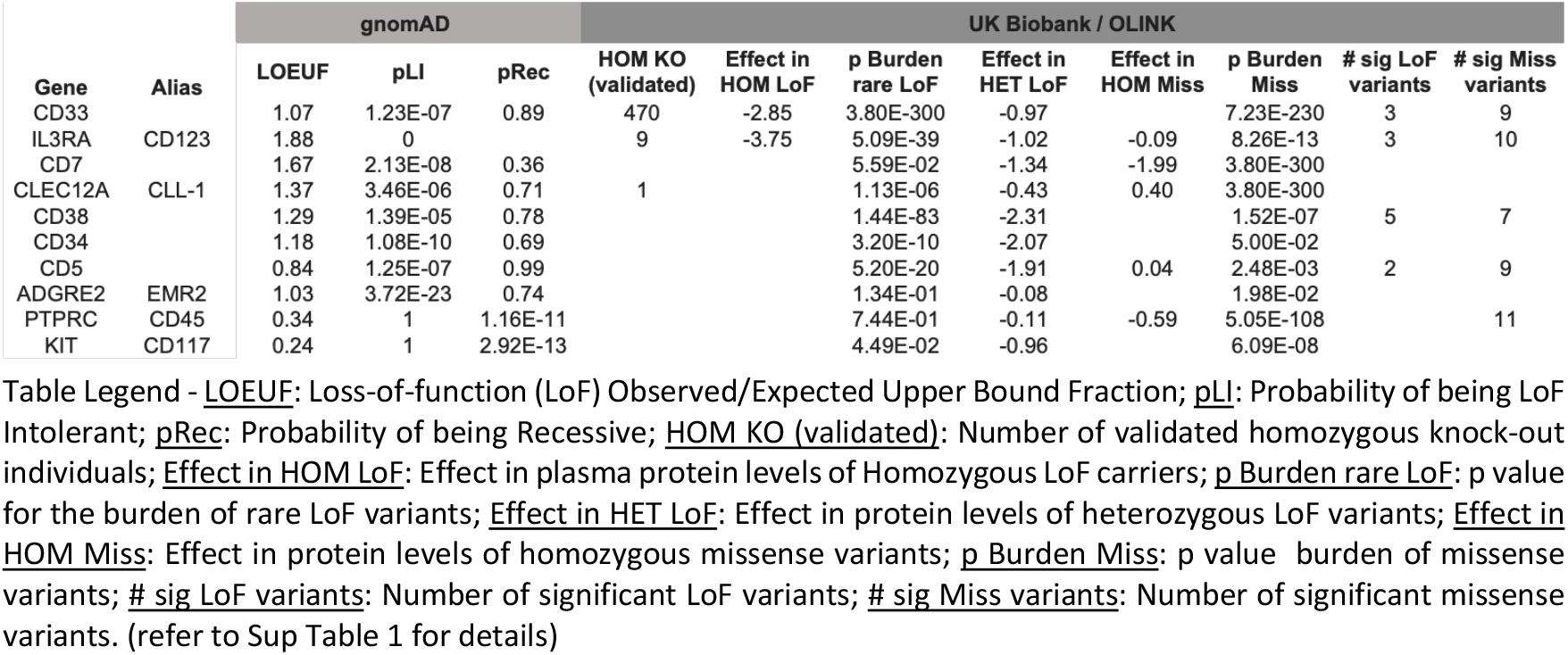
Genetic metrics of candidate hematopoietic genes.

Some antigens, such as *IL3RA* (CD123), display genetic parameters similar to *CD33* with high LOEUF score (> 0.6), low pLI (< 0.9) and homozygous LoF variants validated both genetically and at the protein level in the UKB. Others, such as *KIT* (CD117) resemble *PTPRC* (CD45) and is likely to be functionally important, reflecting its well-established role in hematopoietic stem cell maintenance and development. Genes that fall in the middle of the list such as *CLEC12A, CD38, CD34* are borderline cases requiring larger cohorts and experimental support and functional interrogation to determine dispensability to guide therapeutic targeting. Together, these comparisons demonstrate that our framework discriminates between functionally important and dispensable genes, validating its utility for target prioritization beyond CD33.

### Loss-of-Function Variants in the Human Genome

We extended our analysis to perform an unbiased screen and examine the global distribution of homozygous LoF variants in UKB. Analysis of the UKB exome datasets revealed a rapid increase in identified homozygous LoF variants with larger sample sizes: 1,541 genes in the 200K exome release to 4,186 genes in the 470K release (Figure 2A and 2B). Rare variants (MAF <1%) accounted for the largest increase, demonstrating the value of larger cohorts for capturing signals of dispensability.^20^

**Figure 2:**
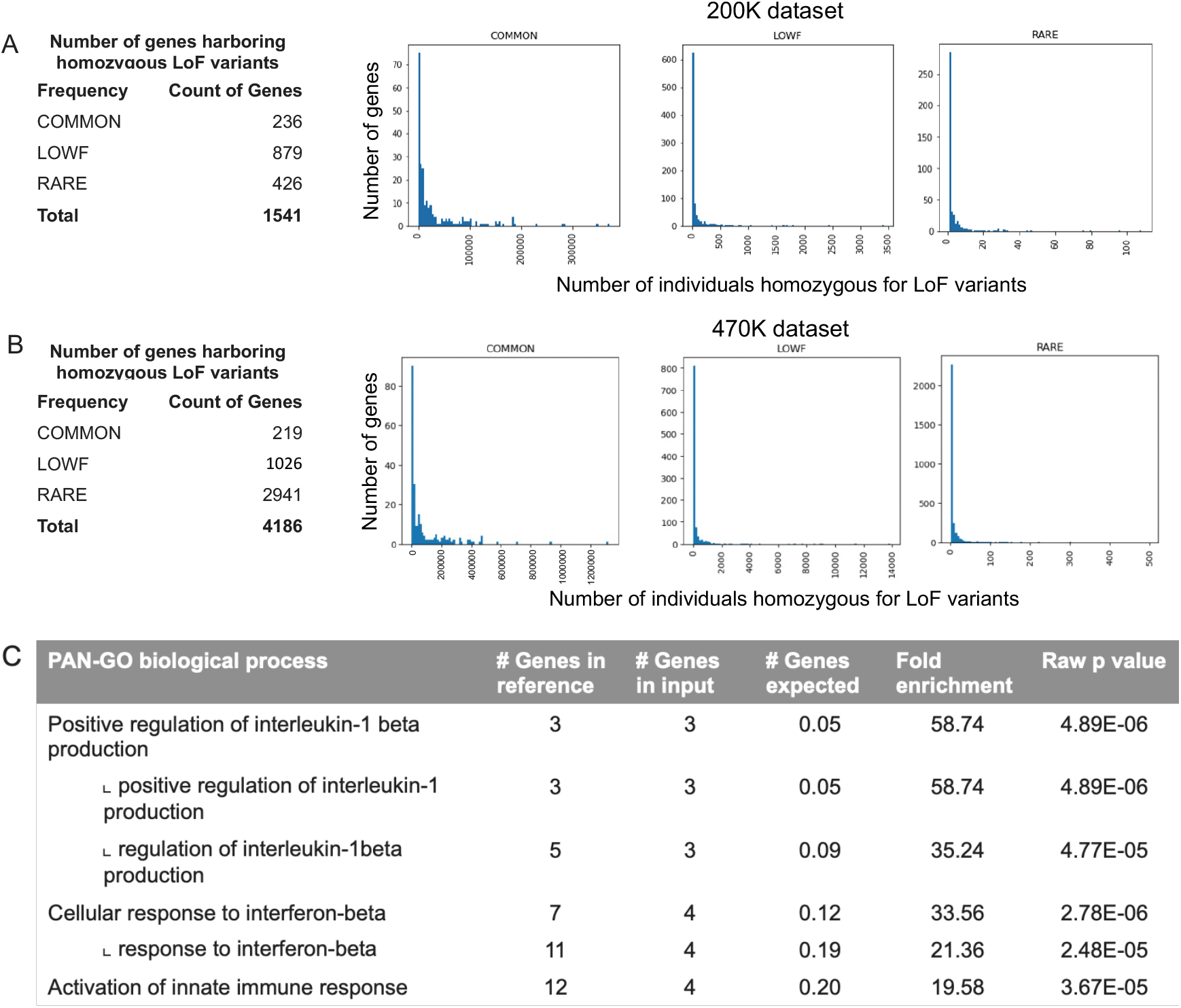
Global analysis of the Loss-of-Function (LoF) variants in the 200K and 470K UK Biobank datasets. A) Distribution of the genes and their frequency in the 200K dataset. B) Distribution of the genes and their frequency in the 470K dataset. C) Gene ontology analysis of hematopoietic genes harboring rare homozygous LoF variants in the 470K dataset. Details of the PAN-GO terms can be found in the methods section.

Given our focus on hematopoietic indications, we next intersected these LoF-tolerant genes with expression data from Tabula Sapiens, restricting to transcripts expressed in at least 1% of bone marrow cells at TPM > 1. These thresholds balance sensitivity (capturing rare but relevant transcripts) with specificity (excluding spurious low-level expression) and are consistent with prior single-cell analyses of hematopoietic compartments. Although the overall proportion of rare LoF variants doubled between releases (27.6% in 200K vs. 70.3% in 470K, Figure 2A and 2B), the proportion of hematopoietic genes harboring homozygous LoF variants observed in both datasets remained the same (11.3% in 200K vs 12.0% in 470K). This stability suggests that most dispensable genes within hematopoietic compartments are already represented at current cohort sizes, supporting a saturation point in the discovery of broadly tolerated LoF events.

We next evaluated whether dispensability signals are enriched within specific biological pathways. Gene ontology analysis of the rare homozygous LoF variant harboring genes from the 470K dataset revealed that genes involved in immune pathways such as interleukin-1 beta production and cellular response to interferon-beta were significantly overrepresented (58.74 fold enrichment, p value 4.89E-06), indicating a greater tolerance for LoF variants in the myeloid lineage of hematopoietic cells (Figure 2C).^21^ These enrichments highlight that LoF tolerance is not uniformly distributed across the genome but tends to occur within immune and cytokine signaling pathways, which may exhibit functional redundancy or compensatory mechanisms, but targeted functional studies will be required to assess this.

### Population-Scale Landscape of Homozygous LoF Variants with Proteomic Readouts

While DNA sequencing predicts which alleles are likely to disrupt gene function, not all annotated LoF variants result in complete inactivation. To distinguish putative from true knockouts, we integrated genotype data with plasma proteomic measurements to validate whether predicted LoF alleles lead to measurable loss of protein. Of 2,958 proteins measured in the UKB Olink plasma proteomics dataset, 467 genes had at least one homozygous LoF carrier; 237 showed (Sup Table 2) concordant reduction in the plasma protein levels in carriers, supporting functional gene inactivation (Figure 3A). Filtering by tissue expression to exclude genes that are expressed in non-hematologic tissues (GTEx, Human Protein Atlas and Tabula Sapiens data) yielded a list of 130 candidate genes (Sup Table 3) with tolerance to inactivation that may be therapeutically relevant for immunotherapy shielding or CAR T enhancement.

**Figure 3:**
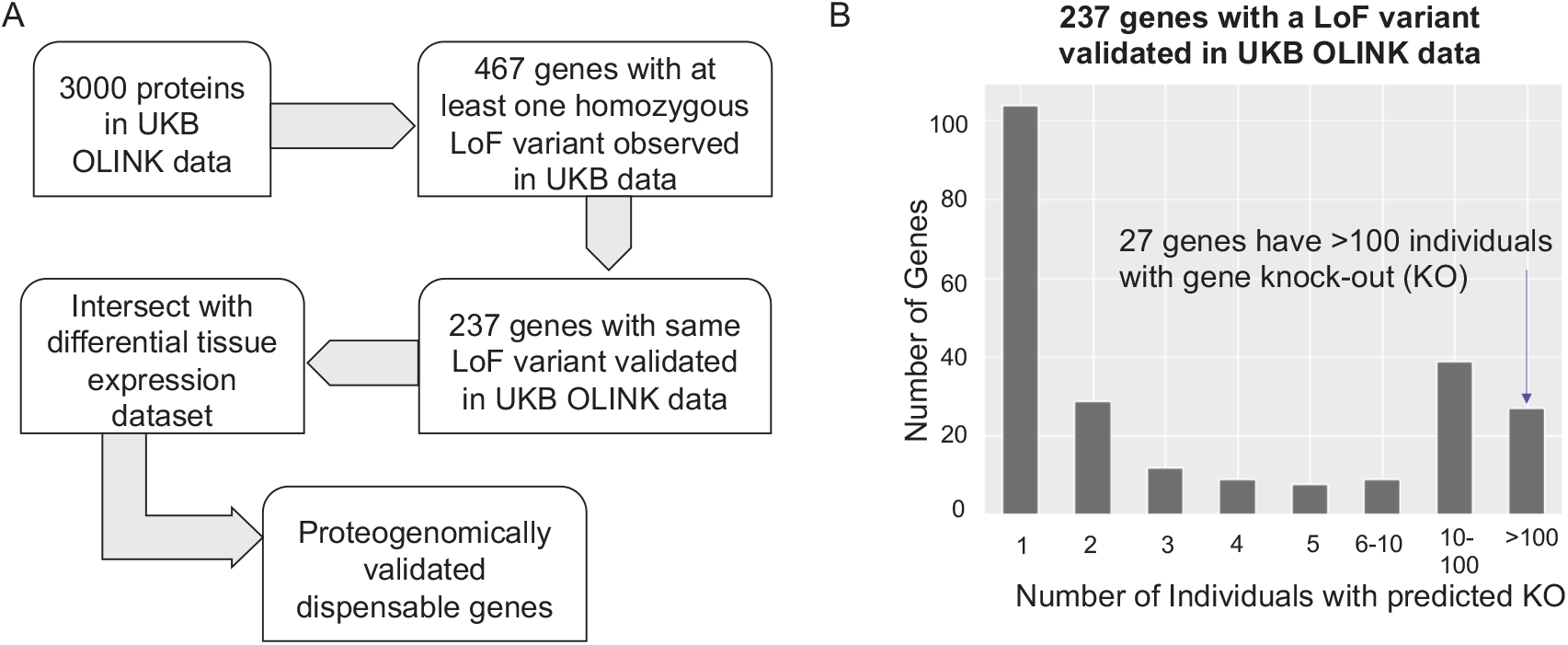
Proteogenomic screen of UK Biobank data to identify dispensable genes. A) Schematic describing the screen methodology. Stepwise filtering pipeline. Genome-wide LoF variant analysis across 470K UKB exomes identified 467 genes with at least one homozygous LoF variant carrier among proteins measured by Olink Explore. Of these, 237 genes showed concordant depletion of the encoded protein, validating functional gene inactivation. Intersection with hematopoietic expression datasets yielded a set of candidate targets, from which the top dispensable genes in blood are highlighted. B) Distribution of the number of homozygous LoF carriers. Among the 237 validated genes, 27 genes had >100 homozygous carriers. Consistent with our operational definition of dispensability, homozygous carriers of validated LoF variants did not exhibit age depletion or clinical phenotypes in UKB, supporting the absence of adverse fitness effects.

### Proteogenomic Evidence Identifies LY75 as a Functionally Dispensable Gene

*LY75* (*CD205*) emerged as the top-ranked candidate from the proteogenomic screen. *LY75* shows minimal evolutionary constraint (LOEUF=0.89, pLI=0) and carries a relatively high frequency of LoF variants (e.g., gnomAD allele frequency of ∼0.0012), including multiple homozygous carriers. Several independent LoF variants, including homozygous carriers (n=2 in 470K dataset), were associated with dose-dependent reductions of circulating LY75 protein levels (Table 2, Figure 4). Across the 470K exome dataset, these analyses included 2 homozygous carriers and 132 heterozygous carriers aggregated across 11 independent LoF variants. In burden tests aggregating high-confidence LoF variants, homozygous carriers showed a mean −4.08 standard deviations (SD) reduction in LY75 while heterozygotes and compound carriers showed intermediate reductions of −2.07 to −2.03 SD (95% CI, excluded 0; Figure 4B).

**Table 2:** Top genes identified in proteogenomic screen.

| Gene | Alias | gnomAD |  |  | UK Biobank / OLINK |  |  |  |  |  |  |  |
| --- | --- | --- | --- | --- | --- | --- | --- | --- | --- | --- | --- | --- |
|  |  | LOEUF | pLI | pRec | HOM KO (validated) | Effect in HOM LoF | p Burden rare LoF | Effect in HET LoF | Effect in HOM Miss | p Burden Miss | # sig LoF variants | # sig Miss variants |
| LY75 | CD205 | 0.89 | 5.27E-48 | 0.99 | 3 | -4.08 | 2.18E-92 | -2.03 | -1.13 | 9.88E-17 | 5 | 29 |
| CD177 | CD177 | 1.56 | 9.35E-14 | 0.12 | 26 |  | 1.37E-41 | -0.87 | 0.26 | 3.01E-113 | 1 | 9 |
| LILRB1 | CD85j | 1.08 | 3.32E-18 | 0.67 | 2 |  | 1.49E-68 | -1.62 | -0.14 | 3.67E-192 | 3 | 14 |
| LILRA2 | CD85h | 1.27 | 1.01E-15 | 0.27 | 260 | -3.47 | 3.80E-300 | -1.77 | -0.25 | 3.80E-300 | 3 | 18 |
| BTN3A2 | CD277 | 1.04 | 3.95E-07 | 0.92 | 8033 | -1.11 | 7.87E-63 | -0.32 | -0.82 | 3.80E-300 | 3 | 9 |
| BTN2A1 | BTN2A1 | 1.46 | 5.38E-16 | 0.09 | 2 |  | 7.52E-17 | -0.96 | 0.15 | 3.80E-300 | 6 | 11 |
| LY9 | CD229 | 1.10 | 6.16E-17 | 0.64 | 1 |  | 4.90E-76 | -2.32 | 0.27 | 2.94E-163 | 7 | 12 |
| LILRA5 | LILRA5 | 1.09 | 2.53E-09 | 0.85 |  |  | 3.21E-11 | -1.73 |  | 2.84E-16 | 4 | 6 |
| CD300LF | CD300LF | 1.11 | 3.47E-08 | 0.86 | 1 | -4.20 | 5.29E-21 | -1.41 | 0.73 | 3.80E-300 | 2 | 12 |
| BST1 | CD157 | 1.64 | 3.16E-18 | 0.02 |  |  | 1.46E-20 | -1.69 | -2.53 | 3.80E-300 | 2 | 3 |
| SIRPB1 | CD172b | 1.19 | 2.02E-08 | 0.76 | 6 | -1.00 | 3.80E-300 | -0.50 | -0.41 | 3.80E-300 | 2 | 18 |
| TLR1 | TLR1 | 1.35 | 7.94E-11 | 0.44 | 1 |  | 6.89E-01 | 0.05 | 0.39 | 3.80E-300 | 2 | 16 |
| CD300C | CD300C | 1.04 | 0 | 0.92 | 14 |  | 5.89E-06 | -0.52 | -1.50 | 1.70E-65 | 1 | 1 |
| FCER2 | CD23 | 1.09 | 1.70E-10 | 0.84 | 1 |  | 1.41E-07 | -1.65 | -0.45 | 4.86E-89 | 2 | 8 |
| TNFRSF13B | CD267 | 1.79 | 4.27E-16 | 0.02 | 1 |  | 2.52E-04 | -0.56 | -0.33 | 2.42E-53 | 1 | 7 |
| CD300A | CD300A | 0.93 | 1.86E-05 | 0.97 | 1 |  | 1.84E-35 | -2.18 | 0.28 | 1.29E-60 | 2 | 7 |

**Figure 4:**
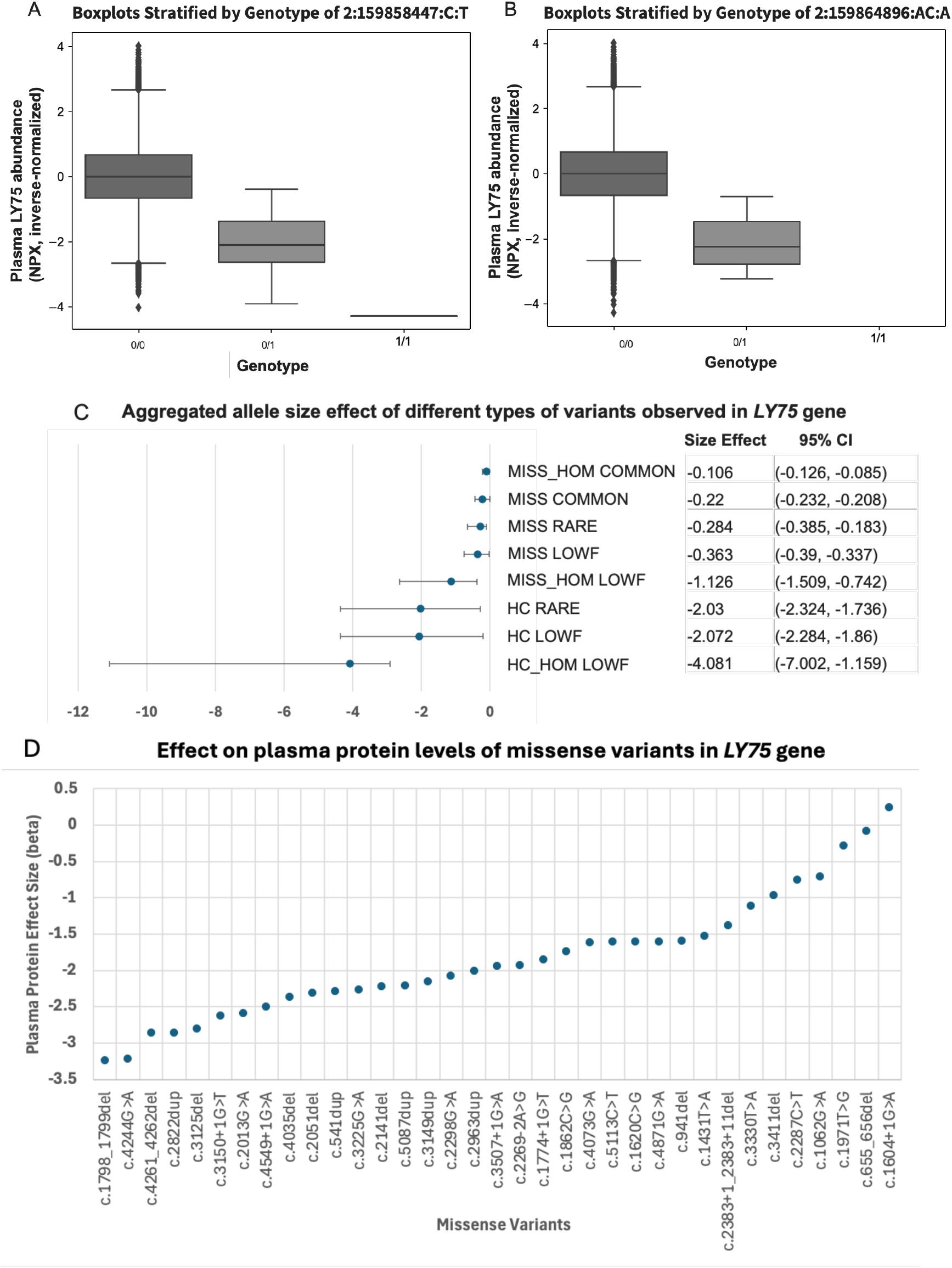
*LY75* LoF variant reduces circulating plasma protein levels. A-B) Plasma LY75 protein abundance in UKB participants stratified by genotype for alleles (A) and (B). Homozygous carriers (1/1; n=2) show the lowest LY75 levels, heterozygotes (0/1; n=132) intermediate, and wild-type (0/0) the highest. C) Aggregated size effect on plasma protein levels for different types of variants observed in the *LY75* gene. MISS_HOM COMMON refers to homozygous missense variants of common frequency, MISS COMMON refers to all missense variants of common frequency, MISS RARE refers to all missense variants of rare frequency, MISS LOWF refers to all missense variants of low frequency, MISS_HOM LOWF refers to homozygous missense variants of low frequency, HC RARE refers to LoF variants of rare frequency, HC LOWF refers to LoF variants of low frequency and HC_HOM LOWF refers to homozygous LoF variants of low frequency. D) Per-variant effect sizes from linear regression analysis (in standard deviations) of individual LoF and missense variants on plasma LY75 protein levels. Most LY75 LoF variants exhibit beta values below −1.0, indicating strong and consistent suppression of LY75 protein expression.

Variant-level analysis corroborated this pattern across the coding sequence. The stop-gain variant p.Trp766Ter exhibited the most extreme effect (β = −2.00, p = 1.5×10^−196^), while most other LoF variants showed consistent suppression (β < −1.0). Several missense variants mirrored this profile, including p.Pro1247Leu (26 homozygotes), which was associated with −1.09 SD reduction in protein abundance (p < 1.0×10^−300^; Figure 4C–D). Consistent with dispensability, no enrichment for adverse clinical phenotypes was observed among homozygous carriers. The consistency and magnitude of these effects provide molecular validation that *LY75* is dispensable and tolerant to loss-of-function.

Although *LY75* encodes an antigen-presenting receptor, no essential hematopoietic functions have been described in human cells.^22^ Because LY75 is an endocytic receptor predominantly expressed on dendritic cells, its inactivation could theoretically affect antigen presentation. However, mouse studies link its deletion to metabolic phenotypes such as obesity, but no defects in hematopoietic stem cell engraftment or multilineage reconstitution have been observed.^23^ Together, these findings provide strong genetic and proteomic evidence that *LY75* is dispensable and nominate it as a strong candidate for therapeutic knockout in donor HSPCs, enabling antigen-directed immunotherapies that spare healthy hematopoietic cells.

## DISCUSSION

Leveraging human genetics databases to de-risk cell and gene therapy targets provides a powerful means to prioritize therapeutic programs and avoid costly drug development failures.^24,25^ Analysis of naturally occurring LoF variants in healthy individuals provides direct evidence of gene dispensability, thereby informing the risks of therapeutic gene editing.^26^ *CD33* provides a validated benchmark of gene dispensability supported by convergent human genetic, experimental, and clinical proof-of-concept.^6-9,27-29^ In the Trem-Cel trial (NCT05945849), CRISPR/Cas9-edited *CD33*-null allografts achieved rapid primary engraftment with durable *CD33*-negative hematopoiesis.^8,9^ Importantly, patients receiving Trem-Cel demonstrated resistance to hematopoietic toxicity during GO maintenance, directly validating the concept that deletion of genetically dispensable antigens can enable safe and effective immunotherapy.^8^ Although CD33 is currently the only dispensable antigen with direct clinical validation, its convergence across human genetics, proteomics, preclinical modeling, and clinical trials makes it a representative benchmark rather than an isolated example. Our framework recapitulates this precedent from population-scale data and extends the same logic to other targets, demonstrating its predictive generalizability.

The list of 237 dispensable candidates provides a rational starting point for therapeutic discovery. These genes can be triaged for experimental validation, used to guide immunotherapy target selection, and re-analyzed as genetic and proteomic datasets expand. In this way, the framework enables a ‘fail fast’ approach that filters unsafe targets before costly preclinical and clinical development.

Extending this approach genome-wide, we identified a set of genes with profiles consistent with dispensability, including *LY75*, which shows strong evidence of tolerance to inactivation. Because the framework is gene-agnostic and built on routinely collected genotype and proteome resources, it can be applied systematically across the genome, re-analyzed as cohorts grow in size and diversity, and extended to future proteomic platforms. While these findings nominate *LY75* as a compelling candidate, functional validation will be vital to confirm dispensability in hematopoietic models and to assess potential immune phenotypes.^22,23^ Accordingly, we view *LY75* not as a validated therapeutic target, but as a high-priority candidate for preclinical testing.

While promising, this framework has limitations. Sex-specific effects were not evaluated given cohort composition (UKB ≈55% female). Current analyses are also constrained by the ancestry composition of UKB, which underrepresents non-European populations, and by proteomic platforms biased toward secreted and extracellular proteins.^30,31,32^ Moreover, the absence of overt phenotypes in UKB participants does not preclude subtle or context-specific effects that may emerge under disease or environmental stress. Dispensability may be tissue- or context-specific, and plasma proteomics may miss critical intracellular functions. Future work will require integrating more diverse cohorts, complementary proteomic technologies, and functional validation in relevant models.

By anchoring target discovery in human data, the approach complements preclinical models and provides a systematic path to de-risk gene-editing targets. *CD33* serves as a benchmark, *LY75* as a novel candidate, and the broader framework as a strategy for refining the therapeutic antigen landscape. Ultimately, integrating human genetics and proteomics offers a rigorous, generalizable, and scalable means to prioritize safe gene-editing targets. As genome and proteome datasets expand across ancestry groups, this framework can be adapted to other tissues, accelerating the rational design of gene-edited therapeutics across disease areas. In doing so, it provides a data-driven complement to empirical discovery pipelines, enabling precision target selection for future CRISPR-based therapeutics.

## ONLINE METHODS

### Samples

Participants were drawn from the UK Biobank, a large prospective cohort of approximately 500,000 UK residents aged 40–69 who were recruited between 2006 and 2010 across 22 assessment centers. Detailed baseline phenotyping—including comprehensive touchscreen questionnaires, anthropometric measurements, and collection of biological samples—was conducted to enable investigation of genetic and environmental determinants of a wide range of health outcomes (Sudlow et al., 2015). All participants provided informed consent under generic research ethics approval (REC reference 11/NW/0382).

## Exome sequencing

Exome sequencing was performed on DNA from 454,787 UK Biobank participants by the UKB Exome Sequencing Consortium. Genomic DNA underwent capture with the IDT xGen Exome Research Panel v1.0 and was sequenced on Illumina NovaSeq 6000 instruments to achieve a mean coverage of ≥20× over targeted coding regions. Sequence reads were aligned to the GRCh38 reference genome and processed through the Regeneron Genetics Center OQFE pipeline for variant calling, after which joint genotyping across all samples produced a population-based VCF containing both single-nucleotide variants and small indels. These cohort-level VCFs enable consistent allele-frequency estimation and large-scale rare-variant association analyses in this ancestrally diverse sample (Backman et al., 2021).

## Proteomics

Proteomic profiling was conducted on plasma samples from 54,219 UK Biobank participants as part of the UK Biobank Pharma Proteomics Project (UKB-PPP). Samples were assayed using the antibody-based Olink Explore 3072 proximity extension assay (PEA), measuring 2,923 unique protein analytes.

https://www.nature.com/articles/s41586-023-06592-6 “Plasma proteomic associations with genetics and health in the UK Biobank | Nature”

This UKB-PPP dataset includes the Normalized Protein eXpression (NPX) values, which represent relative quantification of protein levels on a log2 scale. NPX data has already undergone intra- and inter-assay normalization to account for technical variations and ensure comparability across samples.

Further normalization was required to adjust for potential confounding factors. Rank inverse normal transformation was used to ensure the result was normally distributed. By implementing appropriate normalization procedures, we can reduce the influence of non-biological sources of variation, thereby increasing the reliability and accuracy of the subsequent analyses.

## Protein - QTL analysis

To understand the effect of either single genetic variants or a group of them (burden) we used linear regression models where we adjusted the normalized protein data for relevant covariates including: age, gender, age2, age*gender, age2*gender, 10 principal components, days from collection to Olink analysis and Olink batch to account for potential confounding effects on protein levels. These analyses were performed using hail v 0.2.[ref **Hail Team**. *Hail 0*.*2*. https://github.com/hail-is/hail (accessed July 3, 2025).]

Plots and tables were generated using R v 4.4.0

## PAN-GO analysis

Analysis Type - PANTHER Overrepresentation Test (Released 20200728), is the statistical analysis tool that was selected for the analysis. PAN-GO Version 1.0 Released 2022-03-22 of the PANTHER database, GO database or Reactome pathway database was used for the analysis. Homo sapiens (all genes in database) was used as the reference gene list for the test. The GO biological process complete annotation dataset was used for the analysis using the Fisher’s exact test with FDR correction.

GO:0032731 (positive regulation of interleukin-1 beta production) refers to any process that activates or increases the frequency, rate, or extent of interleukin-1 beta production.

GO:0035458 (cellular response to interferon-beta) refers to any process that results in a change in state or activity of a cell (in terms of movement, secretion, enzyme production, gene expression, etc.) as a result of an interferon-beta stimulus. Interferon-beta is a type I interferon.

GO:0002218 (activation of innate immune response) refers to Any process that initiates an innate immune response. Innate immune responses are defense responses mediated by germline encoded components that directly recognize components of potential pathogens.

## Acknowledgements

We thank Karol Estrada, PhD, for their critical feedback and help with bioinformatics analyses. The authors also acknowledge their colleagues at Vor Bio for their contributions.

## Author contributions

A.D., J.R.L. conceived and designed the study. K.E., A.P., A.D. performed the experiments. Data were analyzed by K.E., A.D. with input from H.G. and J.R.L. The manuscript was drafted by A.D., J.R.L. and all authors critically revised the manuscript for intellectual content. All authors approved the final version of the manuscript.

