## Supplementary Tables for "A Human Genetics Framework for De-risking Gene Editing Targets for Hematopoietic Cell and Gene Therapy"

^1^Vor Bio, Cambridge, MA, 02140, USA.

*Corresponding authors

**Supplementary Data**

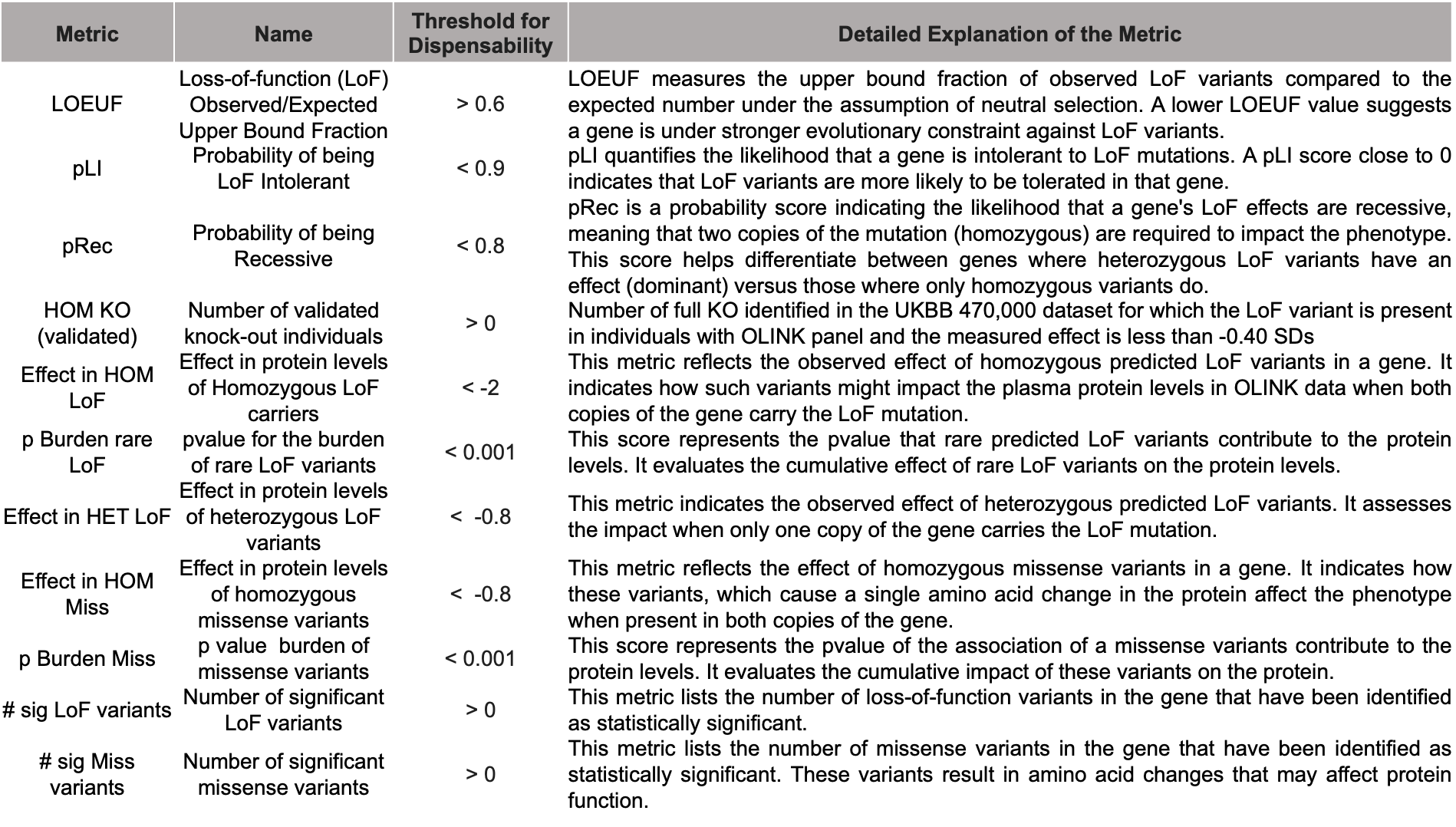

**Supplementary Table 1:** This table provides details of the various metrics and their threshold of dispensability that were utilized to evaluate the dispensability of a gene. Genes with metric values above the threshold of dispensability are more likely to be non-essential and amenable to knock-out.

| ABHD14B | CD177 | CST7 | FOLR3 | KLKB1 | OLFM4 | SLURP1 |
| --- | --- | --- | --- | --- | --- | --- |
| ACADSB | CD1C | CTSE | FTCD | KLRC1 | OTOA | SMPDL3A |
| ADAM8 | CD207 | CTSZ | FUOM | LAIR1 | PADI4 | SORD |
| ADAMTSL4 | CD209 | CXCL11 | GALNT5 | LAIR2 | PCSK9 | SUMF2 |
| ADGRE1 | CD300A | CXCL6 | GIMAP8 | LGALS9 | PDCD1LG2 | TBC1D17 |
| AFM | CD300C | CYB5R2 | GMPR | LILRA2 | PECR | TCN1 |
| AGER | CD300LF | DCBLD2 | GNLY | LILRA5 | PLA2G7 | TGM2 |
| AHNAK2 | CD300LG | DCXR | GP6 | LILRB1 | PON1 | TIMD4 |
| ALDH3A1 | CD33 | DDX58 | GSR | LPA | PON2 | TLR1 |
| AMY2A | CD36 | DFFA | GZMB | LPO | PON3 | TLR4 |
| AMY2B | CD5L | DHRS4L2 | HAVCR1 | LTBP2 | PRTFDC1 | TMPRSS11D |
| ANGPTL3 | CDH17 | DOK2 | HBZ | LY75 | PRTN3 | TMPRSS15 |
| ANXA4 | CDHR5 | DPEP2 | HCLS1 | LY9 | PSAPL1 | TMPRSS5 |
| AP1G2 | CEACAM1 | DPP7 | HGFAC | MAMDC4 | PSCA | TNFRSF10A |
| APOH | CEACAM19 | DRAXIN | HMCN2 | MELTF | PSG1 | TNFRSF10C |
| ART3 | CELA3A | ECHDC3 | HPGDS | MEP1B | PTPRH | TNFRSF13B |
| ASAH2 | CES1 | ECM1 | HPSE | MEPE | PZP | TP53I3 |
| ASPSCR1 | CETN3 | EFCAB2 | HRG | MICALL2 | QPCT | TPMT |
| AZU1 | CFHR2 | ENDOU | HSD17B14 | MINDY1 | RABEPK | TREH |
| BCHE | CFHR4 | ENPP7 | IDI2 | MMP1 | RARRES1 | TRIM58 |
| BGLAP | CFHR5 | ENTPD2 | IDUA | MMP10 | RBKS | TTN |
| BLVRB | CHAC2 | EPPK1 | IFIT3 | MMP12 | REG4 | UPB1 |
| BPIFA2 | CHIT1 | F11 | IL17F | MMP8 | RNASE3 | VAMP5 |
| BST1 | CILP | FCER2 | IL20RA | MMP9 | S100A3 | VIT |
| BTN2A1 | CLEC10A | FCGR2A | IL34 | MNDA | SCGB1A1 | VSIG10L |
| BTN3A2 | CLEC11A | FCGR3B | IL3RA | MPO | SCGB3A1 | VWA1 |
| C1QA | CLEC4A | FCN2 | IL5RA | MRI1 | SCGN | ZBP1 |
| C2 | CLEC7A | FCRL1 | INPP1 | MSR1 | SCLY |  |
| C8B | CNDP1 | FCRL2 | INSL4 | MST1 | SERPINA12 |  |
| C9 | CPA4 | FCRL6 | ITIH1 | MTHFSD | SERPINA9 |  |
| CA1 | CR2 | FCRLB | KIR2DL3 | MUC16 | SERPIND1 |  |
| CAPG | CRACR2A | FGFBP1 | KIR3DL1 | MYOM3 | SHISA5 |  |
| CAT | CRELD1 | FGL1 | KLK12 | NAAA | SIGLEC1 |  |
| CCL26 | CRNN | FKBP7 | KLK13 | NOMO1 | SIRPB1 |  |
| CD109 | CRYGD | FOLR1 | KLK14 | NPC2 | SLAMF8 |  |

**Supplementary Table 2:** This table lists the 237 genes that were observed to be homozygous for knock-out in at least one individual in the UKB genetics data, along with a validated LoF variant in the Olink plasma proteomics data.

| FOLR3 | MSR1 | HCLS1 | MRI1 | BST1 | CHAC2 | TNFRSF13B |
| --- | --- | --- | --- | --- | --- | --- |
| CHIT1 | CAPG | TTN | TLR1 | TBC1D17 | C1QA | FCRL1 |
| SIRPB1 | ABHD14B | ASPSCR1 | FUOM | LGALS9 | CTSZ | TPMT |
| FCRL6 | AZU1 | CRYGD | ADGRE1 | HPSE | MMP9 | FCRL2 |
| BTN3A2 | RBKS | FCGR3B | CST7 | CD300A | CYB5R2 | IFIT3 |
| DHRS4L2 | PZP | CTSE | QPCT | CETN3 | ECHDC3 | CRELD1 |
| CLEC7A | TNFRSF10C | FKBP7 | PECR | LILRA5 | CD300LF | VAMP5 |
| LAIR2 | TRIM58 | PADI4 | AGER | GALNT5 | EFCAB2 | PCSK9 |
| GP6 | NPC2 | GMPR | TP53I3 | SCLY | SHISA5 | AMY2B |
| ZBP1 | CD177 | ACADSB | DPEP2 | ANXA4 | CEACAM1 | CD1C |
| MMP8 | TCN1 | EPPK1 | NOMO1 | BLVRB | SMPDL3A |  |
| LILRA2 | RABEPK | LILRB1 | UPB1 | LY9 | FCER2 |  |
| CD36 | NAAA | ADAM8 | BTN2A1 | TGM2 | GSR |  |
| MAMDC4 | MPO | DOK2 | INPP1 | ADAMTSL4 | CA1 |  |
| OLFM4 | SUMF2 | RNASE3 | PRTN3 | HPGDS | GZMB |  |
| MST1 | CES1 | PON2 | TLR4 | MELTF | MTHFSD |  |
| DPP7 | CLEC11A | CLEC4A | CAT | IDUA | DFFA |  |
| GNLY | SORD | DCXR | KLRC1 | MICALL2 | AP1G2 |  |
| GIMAP8 | CD300C | DRAXIN | IL5RA | CD109 | TNFRSF10A |  |
| PRTFDC1 | FCRLB | MNDA | DCBLD2 | MINDY1 | CRACR2A |  |

**Supplementary Table 3:** This table lists the 130 genes that are expressed in the blood and bone marrow tissue and were observed to be homozygous for knock-out in at least one individual in the UKB genetics data, along with a validated LoF variant in the Olink plasma proteomics data.
